# The ESCRT-III Isoforms CHMP2A And CHMP2B Display Different Effects On Membranes Upon Polymerization

**DOI:** 10.1101/756403

**Authors:** Maryam Alqabandi, Nicola de Franceschi, Nolwenn Miguet, Sourav Maity, Marta Bally, Wouter H. Roos, Winfried Weissenhorn, Patricia Bassereau, Stéphanie Mangenot

**Affiliations:** Laboratoire Physico Chimie Curie, Institut Curie, PSL Research University, CNRS UMR168, 75005 Paris, France; Sorbonne Université, 1 Place Jussieu, 75005 Paris, France; Univ. Grenoble Alpes, CNRS, CEA, Institut de Biologie Structurale (IBS), 38000 Grenoble, France; Moleculaire Biofysica, Zernike Instituut, Rijksuniversiteit Groningen, Nijenborgh 4, 9747 AG Groningen, The Netherlands; Umeå University, Department of Clinical Microbiology & Wallenberg Centre for Molecular Medicine, 90185 Umeå, Sweden

## Abstract

ESCRT-III proteins are involved in many membrane remodeling processes including multivesicular body biogenesis as first discovered in yeast. In humans, CHMP2 exists as two potential isoforms, CHMP2A and CHMP2B, but their physical characteristics have not been compared yet. Here, we use a combination of technics on biomimetic systems and purified proteins to study their affinity and effects on membranes. We establish that CHMP2B binding is enhanced in the presence of PI(4,5)P2 lipids. In contrast, CHMP2A does not display lipid specificity and requires CHMP3 for binding significantly to membranes. On the micrometer scale and at moderate bulk concentrations, CHMP2B forms a reticular structure on membranes whereas CHMP2A (+CHMP3) binds homogeneously. Eventually, CHMP2A and CHMP2B unexpectedly induce different mechanical effects to membranes: CHMP2B strongly rigidifies them while CHMP2A (+CHMP3) has no significant effect. Altogether, we conclude that CHMP2B and CHMP2A cannot be considered as isoforms and might thus contribute differently to membrane remodeling processes.

---

The ESCRT-III (Endosomal Sorting Complex Required for Transport) complex is involved in a variety of cellular contexts^1^ such as biogenesis of multi-vesicular bodies (MVB)^2^, plasma membrane wound repair^3^, neuron pruning^4^, dendritic spine formation^5^, nuclear envelope repair or nuclear envelope sealing during telophase^6, 7^ abscission at a late step of cytokinesis^8, 9^, and budding and release of some enveloped viruses from the plasma membrane of infected cells ^10^. In *Saccharomyces cerevisiae*, the ESCRT-III protein complex comprises four core subunits: Vps20, Vps24, Vps2 and Snf7 (vacuolar sorting proteins 20, 24, 2 and sucrose non-fermenting protein 7), whereas, in *Homo sapiens*, up to 13 proteins form the ESCRT-III family called charged multivesicular body protein (CHMP1-8; IST1) (Supplementary Fig. 1-A). The increased number of ESCRT-III subunits in *Homo sapiens* reflects the functional diversification of the complex in higher organisms ^11^.

In yeast, the sequence of recruitment of ESCRT-III proteins during MVB formation is Vps20 - Snf7 - Vps24 - Vps2, forming a core complex ^12^. Their human homologs are respectively CHMP6 - CHMP4 (A, B, C) - CHMP3 - CHMP2 (A, B). Both CHMP2A and CHMP2B present a high sequence homology with the yeast protein Vps2 and have therefore been considered isoforms. Indeed, CHMP2B appears to be a relatively recent acquisition in the evolution of the ESCRT-III complex resulting from a Vps2 gene duplication event ^11^ (Supplementary Fig. 1-B). Together, CHMP2A and CHMP2B act in most ESCRT-catalyzed membrane remodeling processes, except in MVB formation^5^, where CHMP2A but not CHMP2B is required and neuronal pruning which requires CHMP2B but not CHMP2A. Yet so far, the dual roles of CHMP2A and CHMP2B in the same process remain elusive ^6, 13–17^ (Supplementary Fig. 1-C).

ESCRT-III proteins cycle between an inactive cytosolic state ^18–20^ and an activated lumenal state ^21–23^ leading to filamentous polymers forming spirals or helical tubular structures *in vitro* and *in vivo* ^19, 24–37^. Purified recombinant CHMP2A can coil up into flat snail-like structures ^38^ or form helical tubular polymers with CHMP3 in the absence of membrane ^25^. On the other hand, overexpressed CHMP2B in cells ^32^ leads to the formation of tubular helical structures, but *in vitro* assembly of recombinant CHMP2B has never been visualized, neither alone nor together with CHMP3. SiRNA knockdown of individual ESCRT-III proteins demonstrated a minimal requirement of one CHMP4 and one CHMP2 member for HIV-1 release ^14^. Also CHMP3 acts synergistically with CHMP2A but not CHMP2B ^39^, indicating a distinct role for CHMP2B independently of CHMP3. In contrast, both CHMP2A and CHMP2B are important for cytokinesis ^40^. So far, CHMP2A and CHMP2B have been considered as functional homologs, but practically no study has questioned yet whether CHMP2A and CHMP2B behave similarly upon binding to membranes to validate this hypothesis.

Here we have investigated *in vitro* the functional homology of CHMP2A and CHMP2B in the ESCRT machinery, using biomimetic membrane systems with purified CHMP proteins. We have compared their protein-membrane binding and their mechanical effects on membrane by confocal microscopy, Flow cytometry (FACS), quartz crystal micro-balance with dissipation monitoring (QCM-D) and high-speed atomic force microscopy (HS-AFM). We compare the binding affinities of the proteins to membranes with different charged lipid compositions and investigated the role of CHMP3 on the polymerization of CHMP2A and CHMP2B, respectively. We confirm that CHMP3 works synergistically with CHMP2A for enhancing their mutual binding towards membranes, but reduces the binding of CHMP2B. We establish that CHMP2B binding is enhanced in the presence of PI(4,5)P2 lipids forming a protein network on the membrane surface, whereas CHMP2A+CHMP3 interact via electrostatics with no phosphoinositide binding specificity and bind homogeneously onto membranes. Moreover, we study the mechanical properties of membranes coated with these different ESCRT assemblies. We show by micropipette aspiration, osmotic shock and HS-AFM deformation that CHMP2A and CHMP2B have opposite mechanical effects on the membrane. While CHMP2B highly rigidifies membranes, CHMP2A+CHMP3 have almost no effect on it. Altogether, our study demonstrates that CHMP2A and CHMP2B cannot be considered as functional homologs. Thus, the observed opposite mechanical properties are likely important for understanding the mechanics of membrane remodeling and membrane scission.

## RESULTS

### CHMP2B and CHMP2A display different membrane binding characteristics

Phosphoinositides constitute a minority of the phospholipid family with a concentration lower than 1% in cell membranes. Nevertheless, PIP lipids play an essential role for signaling in cells. PI(3)P is the main phosphatidyl inositide present in the endosomal compartments of the MVB pathway where the ESCRTs were first identified, and this lipid has been used in purified systems to reconstitute MVB formation using yeast proteins ^41^. However, ESCRT-III-mediated processes also occur on membranes enriched in PI(4,5)P2, notably at the plasma membrane, including for instance HIV egress, plasma membrane repair and cytokinesis events, or at the nuclear envelope ^42, 43^. We have first compared the interactions of CHMP2A and CHMP2B with membranes containing different phosphatidyl inositides. To improve protein stability and avoid protein aggregation, CHMP2A was expressed and purified with an MBP tag.

A previous *in vitro* study ^44^ has shown that the interaction of CHMP2B proteins with membrane is significantly enhanced in the presence of PI(4,5)P2 lipids in comparison with DOPS or PI(3,5)P2- membranes. Thus, we compared the preferential binding of CHMP2A versus CHMP2B on GUVS doped with 10% PI(4,5)P2 using confocal imaging.

10% PI(4,5)P2-GUV (see composition 1 in the Methods section) were incubated for 30 min with CHMP2A or CHMP2B proteins at a concentration of 500 nM in the Protein Binding buffer (BP buffer), which has been optimized to ensure the highest protein density on the GUV membrane (Supplementary Fig. 2-A). It has been shown that the displacement or truncation of the C-terminal region of CHMP proteins results in the activation of the protein required for polymerization on membranes ^19, 21, 22, 44^. Thereupon, the constitutively active version of CHMP2A and CHMP2B, i.e. their C-terminal truncated mutants, respectively CHMP2A-ΔC and CHMP2B-ΔC, were used in the following experiments.

While CHMP2B shows a homogenous binding to the GUV in these conditions (Fig. 1-A, first panel), the interaction of MBP-CHMP2A-ΔC is rather weak under the same conditions (Fig. 1-A, third panel). MBP cleavage increases the interaction but also induces aggregation of CHMP2A-ΔC in solution and on the membrane (Supplementary Fig. 2-B).

**Figure 1:**
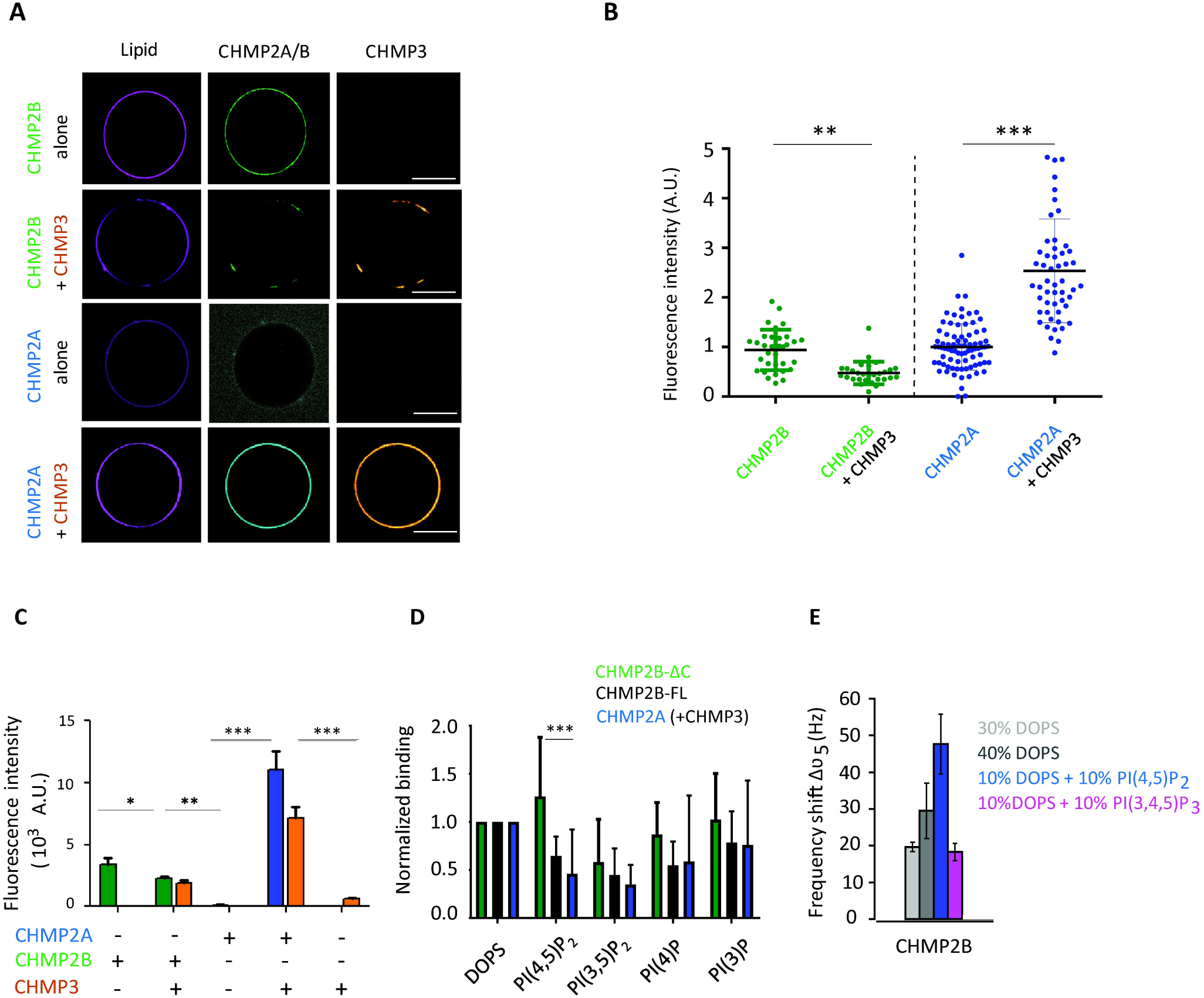
Interaction of CHMP2A versus CHMP2B with charged model membranes. **(A)** Confocal images of 10 % PI(4,5)P2-GUVs incubated with 500 nM CHMP2B-ΔC (first line) (called CHMP2B) and MBP-CHMP2A-ΔC (third line) (called CHMP2A) alone or in combination with 2 μM CHMP3 (second and fourth line respectively). A single confocal plane is shown. Scale bar, 10 µm. Note that in the case of MBP-CHMP2A-ΔC (third line), the laser intensity in the protein channel has been increased (as visible by the higher background intensity) to detect protein binding on the GUV membrane. **(B)** Effect of CHMP3 on MBP-CHMP2A-ΔC and CHMP2B-ΔC binding to 10% PI(4,5)P2-GUVs (same conditions as in Fig. 1A). The fluorescence intensity was measured from the analysis of Spinning Disk microscopy images using the Cell Profiler software. The fluorescence intensity of MBP- CHMP2A-ΔC+CHMP3 and CHMP2B-ΔC+CHMP3-covered vesicles was normalized to the intensity of MBP-CHMP2A-ΔC and CHMP2B-ΔC-covered vesicles, respectively. ***=p-value<0.001; ** = p-value < 0.01 (Student’s t-test). N = 48. **(C)** Quantification by FACS of the fluorescence intensities of MBP-CHMP2A-ΔC ± CHMP3, CHMP2B-ΔC ± CHMP3 and CHMP3 co-polymers bound to 10% PI(4,5)P2-containing GUVs. The concentrations of CHMP2A/B and CHMP3 proteins are, respectively, 500 nM and 2 μM. *=p- value<0.05; ** = p-value < 0.01; ***=p-value<0.001 (Student’s t-test). N = 4 (number of FACS experiment with about 10^4^ counted events per experiment, per condition). **(D)** Quantification of CHMP2B-FL, CHMP2B-ΔC and MBP-CHMP2A-ΔC (CHMP2A) + CHMP3 binding to GUVs containing DOPS and different PIPs by flow cytometry (FACS). Equimolar amount of DOPS and different PIPs (2% mol/mol of total lipids) have been used. Note that data on CHMP2B-ΔC binding to DOPS, PI(4,5)P2 and PI(3,4)P2 were already published in ^44^. Binding efficiencies were normalized to the fluorescence intensity of DOPS-containing vesicles and to the charge of each lipid composition. N = 6 (number of FACS experiment with about 10^4^ counted events per experiment, per condition). **(E)** Resonance frequency shift *Δ𝜗*_5_ in the QCM-D experiments when CHMP2B-ΔC is bound to the different types of supported lipid bilayers (Light grey: 30% DOPS, 70% DOPC; Grey: 40% DOPS, 60% DOPC; Light Blue: 10% DOPS, 10% PI(4,5)P2, 80% DOPC; Magenta: 10% DOPS, 10% PI(3,4,5)P3, 80% DOPC). N = 5.

Previous experiments have shown that in solution, combinations of CHMP2A-ΔC and CHMP3-ΔC as well as CHMP2A-ΔC and CHMP3-FL co-polymerize to form tubular helical structures more efficiently than combinations of CHMP2A-FL and CHMP3-FL or CHMP2A-FL and CHMP3-ΔC ^25^. We have thus tested the effect of CHMP3-FL on the polymerization of CHMP2A-ΔC. In all the following, MBP-CHMP2A-ΔC, CHMP2B-ΔC and CHMP3-FL, will be referred to CHMP2A, CHMP2B and CHMP3, respectively.

After incubation of 10% PI(4,5)P2-GUVs with CHMP2A (or CHMP2B) + CHMP3 (500 nM and 2 μM respectively in BP buffer), we found that CHMP2A strongly binds to GUVs in the presence of CHMP3 (Fig. 1-A fourth panel). The quantification of the fluorescence intensity (see details in Methods section) of CHMP2A on GUVs by confocal microscopy shows that the binding of CHMP2A to the membrane is increased by a factor of at least 2.5 in the presence of CHMP3 (Fig. 1-B), even when the MBP tag is present, justifying that we keep the tag for the rest of our experiments. Surprisingly, when CHMP3 is incubated with CHMP2B, the binding of CHMP2B is no longer homogenous and appears as patches on the GUV colocalizing with CHMP3 (Fig.1-A second panel). The relative amount of CHMP2B on the membrane is decreased by a factor of 2 as compared to the relative CHMP2B amount measured in the absence of CHMP3 (Fig. 1-B).

To quantify the amount of protein bound to the GUV membrane with higher statistics, we have used Flow cytometry (FACS) ^45^ and 2% PI(4,5)P2-GUVs incubated with a combination of CHMP2A, CHMP2B with and without CHMP3, respectively at 500 nM for both CHMP2A and CHMP2B proteins and 2 μM for CHMP3 for 30 min. The fluorescence intensity of the membrane and of the proteins is proportional to the amount of fluorophores in the membrane and proteins bound to it or present in the detection zone, respectively. From the plot of the protein intensity versus lipid signal for all recorded events, we could isolate the signals corresponding to CHMP proteins bound to GUVs and plot the corresponding histogram of these intensities for the different conditions. The median value of this histogram is related to the average density of proteins bound to GUVs. When CHMP2A or CHMP3 are incubated alone with the 2% PI(4,5)P2-GUV suspension, an extremely weak signal is detected by FACS, but as previously observed by confocal microscopy binding increases significantly by almost 100 times-when both proteins are incubated together, in comparison to CHMP2A alone (Fig. 1-C). On the contrary, the presence of CHMP3 decreases the binding efficiency of CHMP2B - by approximately 150 % (Fig. 1-C). We conclude that CHMP2A and CHMP3 synergize in binding to membranes, while CHMP3 acts as a negative regulator for CHMP2B membrane binding *in vitro*.

All the previous experiments were performed with GUVs doped with PI(4,5)P2 lipids. *In vivo*, membranes are enriched with different PIP species depending on their localization. We thus wondered if the behavior of CHMP2A and CHMP2B in the presence or absence of CHMP3 would be affected by the incorporation of other phosphoinositides in the membrane. GUVs were thus produced with 2% of PI(3)P, PI(3,5)P2, PI(4)P or PI(4,5)P2 (see composition 2 in the Methods section), which are enriched at the early endosomes, late endosomes, endoplasmic reticulum/Golgi and plasma membrane, respectively ^42^. They were then incubated with CHMP2A+CHMP3 or CHMP2B alone or in combination with CHMP3 for 30 min to optimize the protein coverage on the membrane. The amount of protein bound to the GUV membrane was analyzed by FACS. The median values of the histograms of binding efficiency for the different PIP species (Supplementary Fig. 2-C) were normalized by the mean value of the distribution of proteins bound to DOPS vesicles (control GUV without PIPs) (Supplementary Fig. 2-D).

Interestingly, experiments performed at a nominal constant phosphoinositide molar fraction in membranes show that the binding efficiency of CHMP2A+CHMP3 does not depend on the PIP species in the membrane. When normalized by the charge of each PIP species, considering that DOPS has a net charge of -1, PI(4,5)P2 (or PI(3,5)P2) of -3 at pH 7.5, and PI(4)P (or PI(3)P) of -2 ^46^, we found that CHMP2A+CHMP3 has almost no preference for any phosphoinositide (Fig. 1-D). Within the error bars, the binding efficiency are more or less equal for all the PIP species tested (i.e. PI(4,5)P2, PI(3,5)P2, PI(3)P and PI(4)P), and lower than to DOPS alone.

In contrast, we found that CHMP2B has a stronger affinity for PI(4,5)P2 than CHMP2A+CHMP3 (Supplementary Fig. 2-D). After charge normalization of the binding density of the PIP species and renormalization by the binding to DOPS lipids, we did not measure a significantly higher binding efficiency of CHMP2B for PI(4,5)P2 membranes than for pure DOPS membranes (Fig. 1-D, p-value = 0.04), nevertheless much stronger than CHMP2A+CHMP3 (Fig. 1-D). Moreover, the binding of CHMP2B is almost doubled on PI(4,5)P2 membranes than on PI(3,5)P2. In contrast, we did not observe such a preference for the full length protein CHMP2B-FL (Fig. 1-D).

FACS experiments suggest a stronger affinity of CHMP2B for PI(4,5)P2 lipids as compared to other phosphoinositides and even for PS lipids. But since the incorporation of phosphoinositides in GUV membrane is in general less efficient as compared to DOPS ^47, 48^, a direct comparison is difficult. Therefore, we used a complementary technique, the Quartz Crystal Microbalance with Dissipation monitoring (QCM-D) to measure CHMP2B binding to supported lipid bilayers (SLB) with a constant net charge fraction (see Methods section). Indeed, the fraction of charged lipids is well preserved during SLB preparation from fusion of small unilamellar vesicles (SUVs) onto solid substrates ^49^. After SLB formation with a defined DOPS or PIP composition (see lipid compositions in the Methods section), CHMP2B proteins were added in the chamber resulting in a shift of the resonance frequency Δϑ_5_ of the quartz sensor, directly related to the amount of protein bound to the surface (Supplementary Fig. 2-E). The amount of proteins adsorbed to the bilayer increased by 50 % when the amount of DOPS was increased from 30 % to 40 % (Fig. 1-E). Indeed, increasing the number of negatively charged lipids in the membrane increases the amount of proteins adsorbed on it. This implies that electrostatic interactions play a key role in mediating the interaction between the proteins and the membrane in agreement with the exposure of basic surfaces in ESCRT-III polymers ^35, 50^. Furthermore, in order to discriminate between the specific affinity for PI(4,5)P2 lipids and electrostatic interactions, we prepared SLBs with a constant total net charge with either 40 % DOPS or 10 % DOPS + 10 % PI(4,5)P2, the total net charge of these SLBs being equivalent. We observe that CHMP2B density is approximately 60 % higher when PI(4,5)P2 lipids are present in comparison with SLBs made of DOPS only (Fig. 1-E). Compared to experiments on GUVs (Fig. 1-D), this higher enhancement is probably due to an effective higher PI(4,5)P2 fraction in the SLBs as compared to the GUVs. Moreover, when PI(4,5)P2 lipids are replaced by the same fraction of PI(3,4,5)P3, the amount of protein bound to the SLB decreases significantly and becomes almost equal to the amount of proteins bound to SLB with 30 % DOPS only, although PI(3,4,5)P3 lipids have a higher negative net charge (-4) as compared to PI(4,5)P2 lipids (-3) ^46, 51^. Altogether, these experiments further support that CHMP2B preferentially interacts with PI(4,5)P2 lipids.

Globally, our results show that while CHMP2B is capable of binding to membrane alone, the binding of CHMP2A to membranes is greatly enhanced by CHMP3 (Figs. 1B, 1C). Additionally, CHMP3 has an opposite effect on CHMP2B and it reduces the membrane its association (Figs. 1B, 1C). Moreover, we found that the binding of CHMP2A+CHMP3 does not depend on the PIP species present in the membrane composition, in strong opposition with the enhanced binding of CHMP2B in the presence of PI(4,5)P2 lipids. This non-specificity of CHMP2A (+CHMP3) proteins to any of the PIP species including PI(4,5)P2 is in agreement with their presence in most cellular processes involving the ESCRT-III complex ^1^, contrary to CHMP2B which is only required for processes occurring at the plasma and nuclear membranes that are enriched in PI(4,5)P2 lipids ^3, 52, 53^.

### CHMP2A and CHMP2B exhibit different supra-molecular assemblies on membranes

Previous studies have shown that cellular overexpression of CHMP2B leads to helical scaffolds deforming the plasma membrane into long rigid tubes protruding out of the cell ^32^. Similarly, CHMP2A + CHMP3 co-assemble in bulk into helical tubes *in vitro* ^25, 39^. Yet, the way CHMP2B or CHMP2A in synergy with CHMP3 assemble *in vitro* onto membranes has not been addressed. The characterization of the effect of these proteins on deformable model membranes is crucial to understand their mechanical properties and their functioning. Hence we studied the supramolecular assemblies of CHMP2B versus CHMP2A+CHMP3 on 10%PI(4,5)P2-GUVs by spinning disk confocal microscopy.

Above 500 nM protein bulk concentration, CHMP2B proteins fully cover the surface of GUVs with no observable distinctive structure, i.e. no inward or outward tubulation (Fig. 2-A first panel). At optical resolution, CHMP2B proteins appear homogeneously distributed on the surface of the vesicles, besides some protein-lipid patches. At bulk concentration lower than 500 nM, CHMP2B proteins form a peculiar reticular-like network wrapping around the whole vesicle (Fig. 2-A second panel). All these observations were performed after 15 min GUV incubation in the protein solution. It suggests that, at high bulk concentration, a reticulum-like network forms transiently, becoming denser with time and leading to an apparent continuous coverage at optical resolution. This CHMP2B network colocalizes with PI(4,5)P2 lipids (Fig. 2-B), indicating that CHMP2B recruits negatively charged PI(4,5)P2 lipids, further confirming the specific interaction between CHMP2B and PI(4,5)P2 lipids.

**Figure 2:**
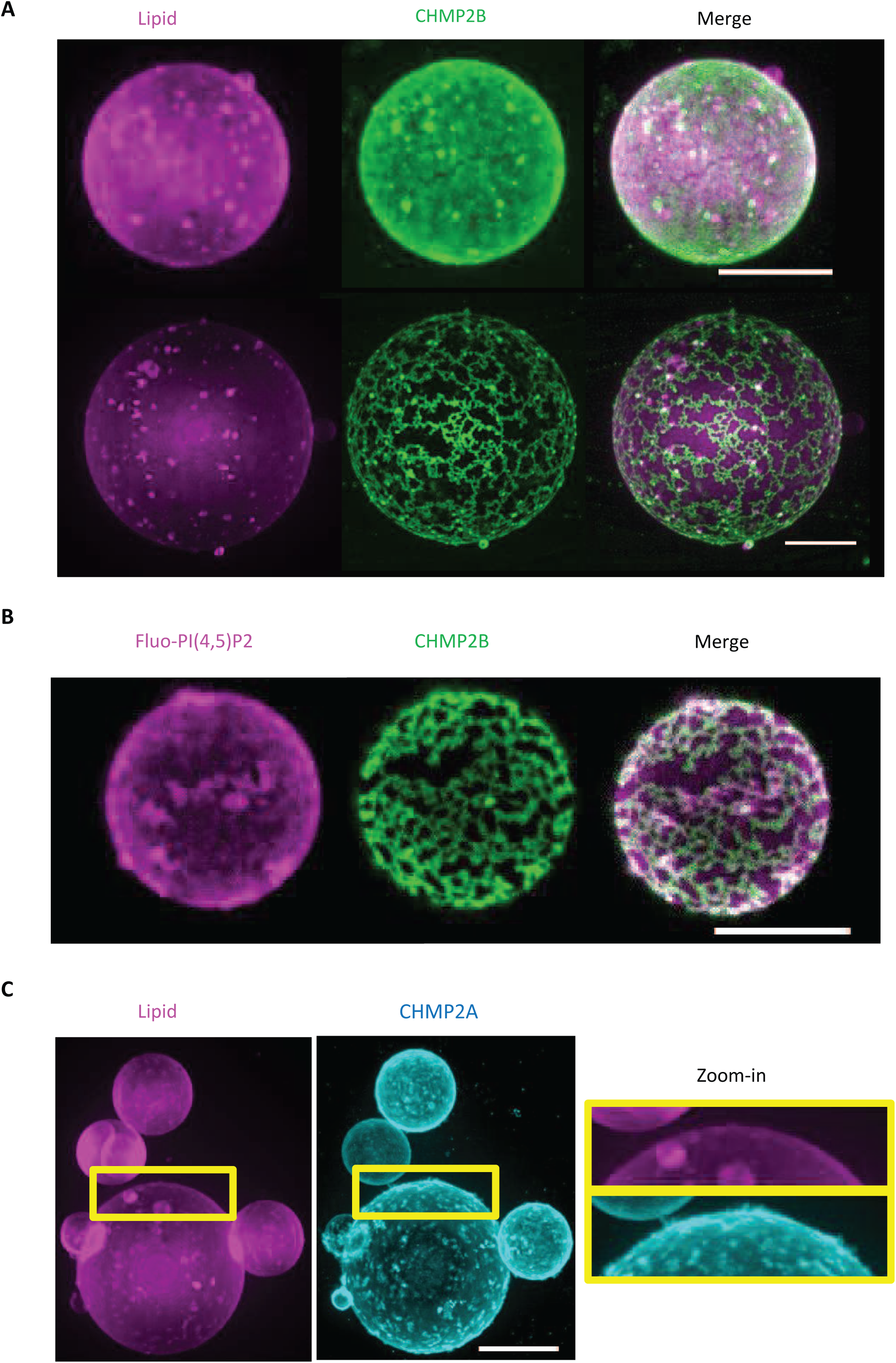
Supramolecular assemblies of CHMP2A + CHMP3 versus CHMP2B on GUV.

**(A)** Spinning disk images of supramolecular assemblies of CHMP2B-ΔC (called CHMP2B) in BP buffer on 10% PI(4,5)P2-GUVs. After 15 min incubation of the GUVs with the protein solution, CHMP2B-ΔC at high bulk protein concentration (1 µM) (first panel) fully covers the vesicle surface, whereas at lower protein concentration (500 nM), CHMP2B-ΔC assembles into a reticular-like network on the GUV (second panel). A z-projection of the whole GUV is shown. Scale bar, 10 µm.

**(B)** Co-localization of Fluo-PI(4,5)P2 and CHMP2B-ΔC on GUVs. A z-projection of the upper part of the GUV is shown. Scale bar, 10 µm.

**(C)** Spinning disk images of supramolecular assemblies of MBP-CHMP2A-ΔC (500 nM) + CHMP3 (2 µM) in BP buffer on 10% PI(4,5)P2-GUVs. MBP-CHMP2A-ΔC (called CHMP2A) fluorescent signal is displayed. After 60 min incubation, the co-polymer covers the vesicle surface in a homogeneous manner with the presence of some protrusions at the surface of the GUV (zoom-in). A z-projection is shown including a zoom-in in the right panel, showing short protrusions at the surface of the GUV. Scale bar, 10 µm.

In contrast, the assembly of CHMP2A+CHMP3 appears very uniform optically, devoid of any visible network, independently of the incubation time and protein concentration (Fig. 2-C and Supplementary Fig. 2-F). In some vesicles (approx. 10 %), we observed CHMP2A (+ non-labeled CHMP3)-containing short out-ward protrusions (Fig. 2-C, and zoom-in). These protrusions were however rarely visible on most of the vesicles. We conclude that CHMP2B and CHMP2A-CHMP3 do not tubulate GUV membranes in this concentration range.

We next investigated whether these proteins perturb the mechanical properties of the membranes.

### CHMP2A and CHMP2B have opposite mechanical effects on model membranes

To study the mechanical effect of CHMP2B and CHMP2A+CHMP3 on membranes, we first used the micropipette aspiration technique developed by E. Evans ^54^, to measure the elasticity of 10 % PI(4,5)P2-GUV (lipid composition 1) coated with CHMP2A or CHMP2B in the presence or not of CHMP3.

In the absence of CHMP proteins, micropipette aspiration of PI(4,5)P2-GUVs easily induced the formation of a characteristic tongue inside the micropipette (Fig. 3-A-first panel). In contrast, PI(4,5)P2-GUVs incubated with a CHMP2B concentration leading to full coverage could not be aspirated and deformed even at high tensions (Fig. 3-A - second panel) (up to 10^−3^ N.m^−1^). However, during aspiration at high tension, in approximately 20 % of the aspirated GUVs (Fig. 3-B), an occasional rupture of CHMP2B protein coat could be observed, allowing the formation of a short tongue devoid of proteins inside the micropipette (Fig. 3-A-third panel). This observation indicates that CHMP2B polymer itself cannot be aspirated or deformed and behaves as a solid shell. Surprisingly, the subsequent CHMP3 incubation with GUVs with pre-formed CHMP2B polymers on their surface resulted in the softening of the CHMP2B shell, which allowed aspiration of the GUV (Fig. 3-A-fourth panel). The quantification of the percentage of aspirated vesicles at a tension of approximately 10^−3^ N.m^−1^ clearly indicates that while less than 20 % of the CHMP2B-coated GUVs could be aspirated in the absence of CHMP3, generally upon shell rupture, almost 100 % of the CHMP2B-coated GUVs could be aspirated when CHMP3 proteins were added (Fig. 3-B). Thus the data suggests that CHMP2B polymers form a rigid shell around the vesicle that cannot be deformed by aspiration even at tensions as high as a few 10^−3^ N.m^−1^ unless CHMP3 is present. The presence of CHMP3 softens this rigid shell allowing its deformation by the micropipette.

**Figure 3:**
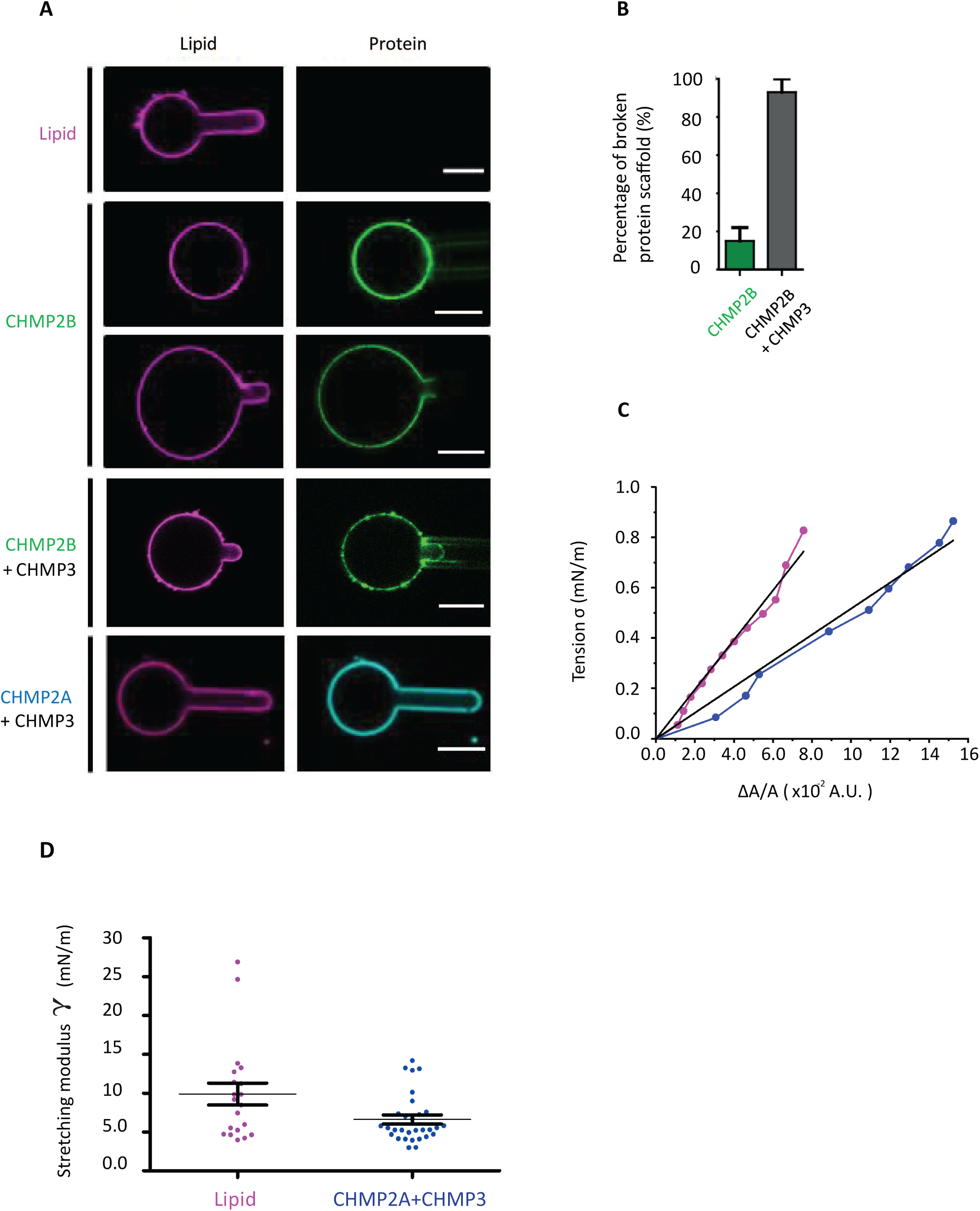
Mechanical properties of GUVs in the presence of CHMP2B versus CHMP2A+CHMP3 measured by micropipette aspiration. **(A)** Representative confocal single-plane images of micropipette aspiration performed at σ ≈ 5.10^−3^ N.m^−1^ on a bare GUV containing 10% PI(4,5)P2-(first panel), and on GUVs coated with CHMP2B-ΔC alone (second panel), CHMP2B-ΔC + CHMP3 (fourth panel) and MBP-CHMP2A-ΔC + CHMP3 (fifth panel) (CHMP2B corresponds to CHMP2B-ΔC and CHMP2A to MBP-CHMP2A-ΔC). The occasional rupture of CHMP2B polymer at high tension (σ ≈ 10^−3^ N.m^−1^) is shown (third panel). Scale bar, 10 µm. N = 30 per condition. **(B)** Percentage of aspirated GUVs at σ ≈ 10^−3^ N.m^−1^ with formation of a tongue inside the micropipette. Comparison between CHMP2B-ΔC-only GUVs and GUVs coated with CHMP2B-ΔC+CHMP3. N = 14 per condition. **(C)** Characteristic curves of the variation of the excess area as a function of the applied tension for a bare GUV (magenta) and a GUV coated with MBP-CHMP2A-ΔC +CHMP3 proteins (blue). The linear fit of each curve is represented (black). **(D)** Box plot of the stretching modulus for bare GUV (N = 20 experiments; magenta) or in the presence of MBP-CHMP2A-ΔC +CHMP3 (N = 30 experiments; blue) interacting with the GUV membrane.

In contrast, PI(4,5)P2-GUVs co-incubated with CHMP2A+CHMP3 could be easily deformed during aspiration with an increase of the tongue length inside the micropipette as a response to the aspiration increase (Fig. 3-A - fifth panel). Fig. 3-C shows the variation of the membrane tension as a function of the fractional excess area, Δα, for two representative experiments. The stretching modulus, χ, calculated from the slope of all the curves for both conditions (see Method section), Fig. 3-D, is within the errors bars not affected by the presence of CHMP2A+CHMP3 on the membrane. It is found to be equal to χ = 10 ± 1 mN.m^−1^ (N = 30 GUVs) for CHMP2A + CHMP3 covered GUVs, similar to the protein-free GUVs, χ = 25 ± 5 mN.m^−1^ (N = 20 GUVs). Note that the value of the stretching modulus for the bare lipid membrane is lower than values reported for dioleoyl, DO, chains, in the presence of cholesterol ^55^, probably because of the absence of a pre-stretching step in our experiments, as usually performed to suppress any pre-existing uncontrolled excess area ^56^. Here, pre-incubation of the GUVs with proteins prevented any pre-stretching of the GUVs in order to limit the contact between pipette and protein-coated GUV and thus adhesion. Nevertheless, our aim was not to measure the absolute stretching modulus of the membranes coated by ESCRTs but to perform measurements relatively to bare lipid membranes in the same experimental conditions. Moreover, the stretching modulus of membrane covered with CHMP2B+CHMP3 could not be measured as the tongue covered with these proteins systematically adhered to the pipette, thus impeded any mechanical measurement. We can however conclude that CHMP2B strongly rigidifies membranes, whereas CHMP2A+CHMP3 membrane interaction does not alter membrane elastic properties.

We next applied different mechanical constraints onto CHMP2B-covered GUVs to test their resistance to mechanical deformations. Spherical GUVs change shape when they are deflated upon an hyperosmotic shock since the surface/volume ratio increases and even becomes unstable when the osmotic shock is too strong ^57^. We thus studied the effect of an hyperosmotic shock on 10% PI(4,5)P2-GUVs fully-covered with CHMP2B by increasing the osmolarity in the external medium by salt or sugar addition. We carefully checked that the buffer change did not induce unbinding of the ESCRT-III proteins from the membrane. An osmotic shock equal to 150 % (osmolarity of the external medium ≥ 190 mOsm.L^−1^) transforms spherical protein-free GUVs into elliptical vesicles (Fig. 4-A) with an average eccentricity index equal to 0.72 ± 0.11 (Fig. 4-B) (note that the eccentricity index is the ratio between-foci distance and the major axis-length of an ellipse. It ranges between 0 (for a circle) and 1 (for a linear segment)). At higher osmotic shock, GUVs were completely destabilized in the absence of proteins. On the other hand, CHMP2B-covered GUVs better preserved their spherical shape for the same 150 % osmotic shock (Fig. 4-A) with an average eccentricity index equal to 0.35 ± 0.03 (Fig. 4-B). Moreover, in contrast with bare membranes, vesicles covered with CHMP2B proteins could even stand a 300 % osmotic shock in a solution at 500 mOsm.L^−1^, showing again that CHMP2B polymer assembly on GUVs’ surface preserves vesicles from deformation by forming a rigid shell.

**Figure 4:**
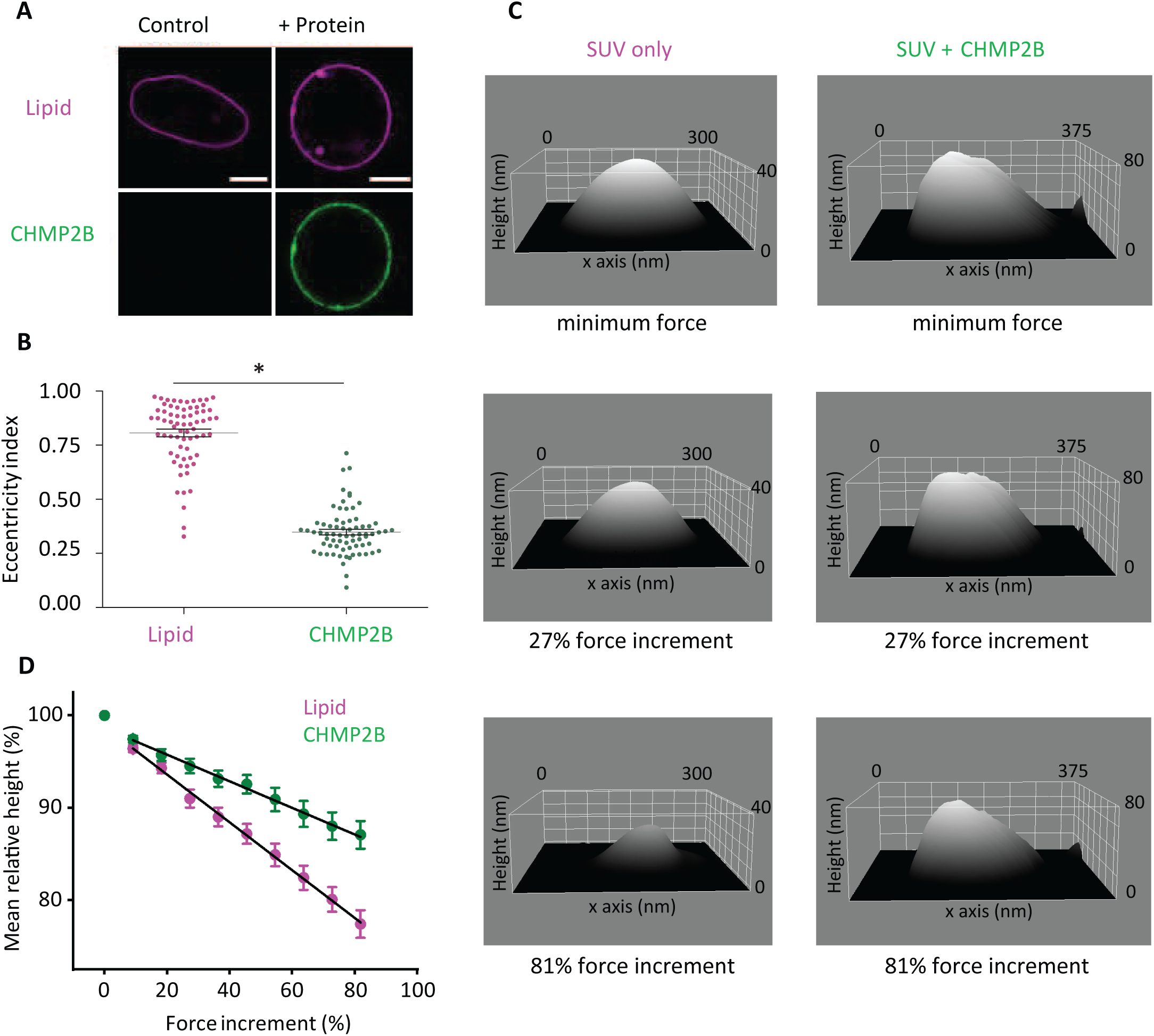
Study of CHMP2B mechanical properties by osmotic shock and HS-AFM indentation. **(A)** Confocal images of GUV submitted to an osmotic pressure difference equal to 150% (Osmolarity inside and outside the GUV are respectively 120 mOsm.L^−1^ and 315 mOsm.L^−1^) without (top) or with (bottom) pre-incubation with CHMP2B-ΔC (noted CHMP2B) at a concentration of 1 μM. Scale bar, 10 µm. **(B)** Eccentricity index of GUVs (alone or covered with CHMP2B-ΔC polymer) pre-formed in a solution with an osmolarity of 120 mOsm.L^−1^ and transferred to a hypertonic solution with an osmolarity of 315 mOsm.L^−1^ (relative osmotic pressure = 150%). *=p-value<0.05 (Student’s t- test). N = 40. **(C)** HS-AFM images of a bare vesicle (left) and a vesicle covered with CHMP2B-ΔC proteins (right). The vesicles with CHMP2B show an increase in surface roughness with respect to the vesicles without CHMP2B (first panel). The deformability of CHMP2B-coated SUVs upon increased applied force is shown at intermediate force increments of 27% (second panel) and at higher force increment, 81 % (third panel). **(D)** Decrease in relative height of bare vesicles (purple) and vesicles coated with CHMP2B-ΔC (green) as a function of force increment. 100% height corresponds to the initial height value, and 0 % force increment corresponds to the minimal imaging force. The inverse of the slope of these graphs is directly related to the stiffness k of the vesicles. The error bar represents the standard error of the mean (SEM).

We next aimed to determine the effect of CHMP2B on the mechanical properties of membranes at the nano-scale, mimicking the relevant cellular dimensions. To do this we applied a High Speed-AFM imaging based deformation approach using Small Unilamellar Vesicles (SUVs) with a typical diameter between 20-80 nm. First of all, a difference in surface roughness is observed between the two kinds of vesicles (with and without CHMP2B). Bare SUVs show a smooth surface and CHMP2B-coated vesicles possess a rougher surface, indicating the presence of the protein on the outside of the vesicles (Fig. 4-C first panel). Next, we increased the imaging force and it can be observed that the vesicles are progressively more deformed. The deformation of the SUVs is measured by recording the height change. (Fig. 4-C second and third panels). To assure that the vesicles had undergone elastic deformation, even at the maximum applied imaging force, the imaging force was reduced again to the minimum value at the end of the experiments. Only vesicles that bounced back to more than 90% of the initial height were considered for the analysis and typically the vesicles did recover their shape and size (movie S1). After assuring that the vesicles behave elastically, we deduce the relative stiffness *k_rel_* of the vesicles from the inverse of the slope of height change versus force increment (Fig. 4-D). In Supplementary Fig.3 we show all the data points on both types of vesicles and the transformation from absolute height to relative height. It can be observed that there is a clear difference between the two kinds of vesicles. For bare SUVs we find *k_rel_* ≈ 3.7 ± 0.1 (N=31, mean ± SD) and for CHMP2B-coated vesicles we find a 2-fold stiffening with *k_rel_* ≈6.7 ± 0.1 (N=32, mean ± SD) revealing that CHMP2B also stiffens membranes at physiologically relevant length scales.

## DISCUSSION

The objective of our study was to compare the membrane binding properties of ESCRT-III proteins CHMP2A and CHMP2B *in vitro* in order to determine their capacity to substitute each other during membrane remodeling processes.

First, we show that CHMP2A binding is strongly enhanced in the presence of CHMP3 to bind membranes, in agreement with previous *in vivo* and *in vitro* studies^25, 26, 39, 58^, whereas CHMP2B interacts with membrane independently of CHMP3. This is in agreement with the synergy exerted by CHMP3 in the presence of CHMP2A and with the absence of synergy exerted by CHMP3 in the presence of CHMP2B on HIV-1 budding ^25, 26, 39, 58^. In fact, we show further that CHMP3 acts as a negative regulator of CHMP2B for membrane interaction and polymerization, as indicated by the reduced binding of CHMP2B to GUV membranes in the presence of CHMP3.

Second, we confirm that CHMP2B displays a stronger binding for PI(4,5)P2 containing membranes as compared to other phosphoinositides and DOPS lipids ^44^. In contrast CHMP2A and CHMP3 require only negatively charged membranes for binding with no preference for specific lipid head groups. The binding affinity with PI(4,5)P2 lipids is in agreement with the spontaneous localization of CHMP2B to the plasma membrane enriched in PI(4,5)P2 ^42^ upon VPS4 knockdown ^32^. In this context, all ESCRT-driven remodeling processes that involve CHMP2B, take place at PI(4,5)P2-containing membranes such as HIV1 budding, plasma membrane repair, cytokinesis nuclear envelope reformation and dendritic spine formation ^8, 43^. Thus our *in vitro* data suggest that CHMP2B recruitment to membranes may be regulated by PI(4,5)P2 and thus PIP signaling.

While CHMP2A and CHMP3 assemble homogenously on the GUV membrane, CHMP2B forms a striking reticulum-like structure at the GUV surface at low density. The network colocalizes with PI(4,5)P2 indicating clustering of PI(4,5)P2 upon CHMP2B network formation. This network formation leads to a strong mechanical stiffening of the membrane. The CHMP2B coat behaves as a rigid shell that can be occasionally fractured upon strong micropipette aspiration. The effect of CHMP3 on CHMP2B membrane binding/polymerization influences also the stiffness of the membrane by softening it compared to CHMP2B only coated membranes. At smaller scale, on SUVs, this stiffening is also observed. In contrast, the mechanics of GUVs coated with CHMP2A + CHMP3 is almost unchanged. Previous experiments performed on yeast ESCRT-III proteins reported a plastic deformation of membrane coated with Snf7 ^30^. As a consequence, the mode of action of the ESCRT-III may be regulated by the balance of stiffening and elastic behavior.

In general, in the concentration range explored in our study, with both CHMP2A (+CHMP3) and CHMP2B, we did not observe spontaneous GUV membrane tubulation. Tubulation depends on protein spontaneous curvature, surface fraction, membrane tension, protein-protein interactions and protein assembly stiffness ^59^. Considering the propensity of the ESCRT-III proteins to form spiral or helical polymers in solution, we could have expected that they might also induce membrane deformation upon polymerization on a lipid membrane. One possible explanation is that we have not included CHMP4, an ESCRT-III member essential for all ESCRT-catalyzed processes ^8^. Although CHMP4 assembles on flat membranes ^30, 60^, it seems to prefer negative membrane curvature for interaction ^61^. Thus, the CHMP2B and CHMP2A+CHMP3 membrane binding observed here has produced assemblies that are different from ESCRT-III assemblies observed *in vitro* ^62, 63^ lacking spontaneous curvature and/or being too elastic to deform membranes.

The differences observed between CHMP2A and CHMP2B with regard to their membrane interaction and their capacity to affect membrane rigidity, indicate that both isoforms exert different functions that require different mechanical properties during ESCRT-catalyzed membrane remodeling processes. As an example, the CHMP4B isoform is likely present in the ESCRT-III spirals formed at the mid-body during cytokinesis ^64, 65^ whereas CHMP4C is implicated in abscission control ^15^. The increased rigidity imposed by the CHMP2B network might be important for dendritic spine maintenance ^66^ where it might limit protein diffusion, in agreement with experiments showing that CHMP2B forms a diffusion barrier at membrane necks reconstituted *in vitro* ^44^. It might also significantly contribute to the mechanical property of the ESCRT-III spirals at the cytokinetic bridge that become very loose when CHMP2B is depleted ^65^. Interestingly, CHMP2B function might be modulated by CHMP3, which limits CHMP2B-membrane interaction and softens the CHMP2B assembly. This indicates that *in vivo* CHMP3 either limits CHMP2B polymerization or/and copolymerizes with CHMP2B into a structure with different mechanical properties, in agreement with observations of copolymerization of ESCRT-III proteins in solution ^60^.

We thus propose that CHMP3 could play a key regulatory role in the sequence of recruitment of CHMP2B and CHMP2A and in their respective stoichiometry on the membranes during ESCRT-III function. In late steps of cytokinesis, pulling forces exerted by daughter cells on the intercellular bridge appear to regulate abscission, allowing daughter cells to remain connected until they have settled in their final locations. Moreover, counter-intuitively, a release of tension conducts membrane scission ^67^. Thus, membrane unaltered softness may be important at the very last stage of the membrane scission event carried out by the ESCRT-III complex, whereas a rigid structure would oppose this process. However, a certain degree of membrane rigidity might help the constriction process prior to scission, but at this stage, it is difficult to conclude on this aspect.

In summary our data provides evidence that CHMP2A and CHMP2B polymerize differently on membranes and thereby impose different mechanical properties on the membrane structure. Our data thus strongly argue against a sole redundancy of the CHMP2A and CHMP2B proteins and indicate that different isoforms exert complementary functions within the ESCRT-III system.

## METHODS

### Reagents

DOPC (1,2-dioleoyl-sn-glycero-3-phosphatidylcholine), DOPS (1,2-dioleoyl-sn-glycero-3-phospho-L-serine), DOPE (1,2-dioleoyl-sn-glycero-3-phosphatidylethanol-amine), cholesterol (cholest-5-en-3ß-ol), PI(3)P (1,2-dioleoyl-sn-glycero-3-phospho-(1’-myo-inositol-3’-phosphate)), PI(3,5)P2 (1,2-dioleoyl-sn-glycero-3-phospho-(1’-myo-inositol-3’,5’-bisphosphate)), PI(4)P (L-α-phosphatidylinositol-4-phosphate), PI(4,5)P2 (L-α-phosphatidylinositol-4,5-bisphosphate), BODIPY TMR-PtdIns(4,5)P2, C16 (red PI(4,5)P_2_), 1-oleoyl-2-6-[4-(dipyrrometheneboron difluoride) butanoyl] amino hexanoyl-sn-glycero-3-phosphoinositol-4,5-bisphosphate (TopFluor PI(4,5)P2), and Egg Rhod PE (L-α-phosphatidylethanolamine-N-lissamine rhodamine B sulfonyl) were purchased from Avanti Polar Lipids, Inc (Avanti Polar Lipids, U.S.A.). Stock solutions of lipids were solubilized in chloroform at a concentration of 10 mg.mL^−1^, except for cholesterol and Egg Rho PE dissolved respectively at a concentration of 20 mg.mL^−1^, and 0.5 mg.mL^−1^ and PIPs, which were solubilized in a mixture of chloroform/methanol (70:30) (v/v) at a concentration of 1 mg.mL^−1^. All stock solutions were kept under argon and stored at −20°C in amber vials (Sigma-Aldrich, France).

### Expression, purification and labelling of proteins

Expression and purification of MBP-CHMP2A-ΔC (residues 9-161) and CHMP3-FL (residues 1-122) was performed as described in ^18^. A final gel filtration chromatography step on a superdex200 column was performed in a buffer containing 20 mM Hepes pH 7.6, NaCl 150 mM.

CHMP2B-FL (residues 1-222) and CHMP2B-ΔC (residues 1-161) were expressed and purified as previously described ^32^. Both constructs contain a C-terminal SGSC linker for cysteine-specific labelling. Cells were lysed by sonication in 50 mM Tris pH 7.4, 1M NaCl, 10mM DTT, complete EDTA free and the soluble fraction was discarded after centrifugation. The pellet was washed three times a buffer containing 50 mM Tris pH 7.4, 2M UREA, 2% TritonX100, 2mM β-mercaptoethanol and a final wash in 50 mM Tris pH 7.4, 2mM β-mercaptoethanol. CHMP2B (-FL and -ΔC) was extracted from the pellet using a buffer composed of 50 mM Tris pH7.4, 8M guanidine, 2 mM β-mercaptoethanol over night at 4°C. Further purification of solubilized CHMP2B included Ni^2+^-chromatography in 50 mM Tris pH7.4, 8M urea, refolding by rapid dilution into a buffer containing 50 mM Tris pH7.4, 200 mM NaCl, 2 mM DTT, 50 mM L-glutamate, 50 mM L-arginine at a final concentration of 2 μM. Refolded CHMP2B was concentrated by Ni^2+^ chromatography in a buffer containing 50 mM Tris pH7.4, 200 mM NaCl. A final gel filtration chromatography step was performed on a superdex75 column in the buffer containing 50 mM Tris pH 7.4, 100 mM NaCl.

Following expression, CHMP proteins were concentrated, labelled overnight at 4°C with a ratio of Alexa labelling dye per protein of 2 to 1. MBP-CHMP2A-ΔC, CHMP3-FL and CHMP2B (-ΔC and -FL) were labelled with Alexa 488 succimidyl ester, Alexa 633 succimidyl ester and Alexa 488 C5 maleimide (Thermo Fisher Scientific) respectively. The excess of free dyes was removed by salt exchange chromatography. Immediately after labelling, all aliquots were frozen in liquid nitrogen with 0.1% of methyl cellulose (Sigma Aldrich) as cryoprotectant. All aliquots were kept at -80°C prior to experiments.

### GUV preparation for confocal, Spinning Disk and FACS experiments

GUVs were prepared by spontaneous swelling on polyvinyl alcohol (PVA)-based gels ^68^. A thin lipid solution is deposited on a PVA gel (5% PVA, 50 mM Sucrose, 25 mM NaCl and 25 mM Tris, at pH 7.5), dried under vacuum for 20 min at room temperature and rehydrated with the growth buffer at room temperature. Vesicles form within 45 min and are extracted by pipetting directly from the slides on top of the PVA gel.

#### Composition 1

For confocal and Spinning Disk microscopy experiments, lipid stock solutions were mixed to obtain DOPC/DOPS/DOPE/Cholesterol/PI(4,5)P2/PE-Rhodamine (54.2:10:10:15:10:0.8) (molar ratio) at a concentration of 3 mg.mL^−1^ in chloroform.. In the following, this GUV composition will be referred to as 10 % PIP2-GUV. In order to detect the PI(4,5)P2 lipid signal, PE-Rhodamine in the PIP2-GUV lipid stock solution was replaced by TopFluor PI(4,5)P_2_ with a molar ratio of PI(4,5)P2/TopFluorPI(4,5)P2 of (8:0.5) referred to as FluoPIP2-GUV.

#### Composition 2

For FACS microscopy experiments, lipid stock solutions were mixed to obtain DOPC/DOPE/Cholesterol/PI(4,5)P2/PE-Rhodamine (72.2:10:15:2:0.8) (molar ratio) at a concentration of 3 mg.mL^−1^ in chloroform. In the following, this GUV composition will be referred to as 2% PIP2-GUV. To compare CHMP protein binding to different PIP species, we replaced PI(4,5)P2 lipids at equal molar ratio by PI(3)P, PI(4)P and PI(3,5)P2 lipids, respectively. In the following, these GUV compositions will be referred to as 2% PI(3)P-GUV, 2% PI(4)P-GUV and 2% PI(3,5)P2-GUV.

### SUV preparation for QCM and AFM experiments

After preparation of lipid <u>Composition 1</u>, at 3 mg.mL^−1^, in chloroform the solvent was evaporated by rotating the vial under a gentle stream of nitrogen, at room temperature and then was placed under vacuum for 20 min at room temperature. The dried lipid film was rehydrated in the appropriate growth buffer solution to obtain a final concentration of 1 mg.mL^−1^. The solution was vortexed for 2 min and then extruded 11 times through a polycarbonate track-etched membrane with pore sizes of 100 nm ^69^ or sonicated for 5 min until obtaining a clear colorless solution for Small Unilamellar Vesicles (SUV) formation. Produced SUVs were either used freshly for QCM-D experiments and for HS-AFM indentation experiments or stored at -20°C in amber vials (Sigma-Aldrich, France) for further use. In the following, this SUV composition will be referred to as 10 % PIP2-SUV.

To compare CHMP2B protein binding in the absence of PI(4,5)P2 and to increase the net negative charge of the membrane for QCM-D experiments, SUVs were produced containing DOPC/DOPS/DOPE/Cholesterol/PE-Rhodamine (44.2:30:10:15:0.8) (molar ratio) or (34.2:40:10:15:0.8), at a concentration of 3 mg.mL^−1^ in chloroform referred to as 30% DOPS-SUV and 40% DOPS-SUV, respectively. Moreover, to compare CHMP2B protein binding to a membrane incorporating a higher amount of negative charges as well as PIP lipids, we replaced the 10% molar ratio of PI(4,5)P2 in the PIP2-SUV by 10% molar ratio of PI(3,4,5)P3 lipids. In all QCM-D experiments, quartz crystal resonance frequency shifts were measured at the overtone 5 of the oscillating crystal and therefore defined as Δϑ_5._

### CHMP supramolecular assembly on GUVs observed by fluorescence microscopy

Freshly produced 10% PIP2-GUVs were incubated with CHMP proteins at concentrations ranging from 50 nM to 2 µM in BP buffer (Tris 25 Mm, NaCl 50 mM pH 7.5) in isotonic conditions for 15 to 30 min. Then, CHMP-coated GUVs were diluted 20 times and transferred to the observation chamber, previously passivated with the β-casein solution and rinsed twice with BP buffer. Supramolecular assembly of CHMP proteins on GUVs was visualized on an inverted Spinning Disk Confocal Roper/Nikon. The spinning disk is equipped with the camera, EMCCD 512x512 Andor Technology (pixel size:16 µm), an objective (100x CFI Plan Apo VCoil NA 1,4 WD 0,13) and 3 lasers (491, 561, 633 nm 100 mW). The exposure time for all images was 50 milliseconds.

To further characterize and compare the interaction of CHMP proteins on GUVs, we measured total intensity of the protein on the vesicle and normalized this value by the GUV area. Image acquisition for protein quantification was performed using a confocal microscope composed of an inverted microscope (Eclipse TE2000 from Nikon), two objectives (60x water immersion and 100x oil immersion), a C1 confocal head from Nikon, three lasers (λ=488 nm, λ=561 nm and λ = 633 nm). One confocal plane image was taken for each set tension.

### FACS experiment for protein-lipid binding assay

2 % PI-GUVs and CHMPs fluorescence intensity was measured with a BD LSRFORTESSA flow cytometry instrument. Data analysis was performed with BD FACS Diva software.

The collected GUVs were transferred in BP buffer and incubated 30 min with CHMP proteins at 500 nM. The vesicle concentration was adjusted in order to count about 10000 events per condition every 60 seconds at high speed.

2 % PI-GUVs were labeled with Egg Rhod PE (0.8 % w/w), CHMP2B labelled with Alexa 488, CHMP2A labeled with Alexa 488 and CHMP3 labeled with Alexa 633. Alexa 488, was excited with a 488-nm laser, and the emission was detected through a 530/30 standard bandpass filter. Alexa 633, was excited with a 633-nm laser, and the emission was detected through a 670/30 bandpass filter. Egg Rhod PE, was excited with a 532-nm laser, and the emission was detected through a 610/20 bandpass filter. Two signals were closely analyzed: the protein fluorescent signal and the lipid fluorescent signal. Thus the fluorescence intensity of the membrane and the fluorescence intensity of the proteins are respectively proportional to the amount of fluorophores in the vesicle and proteins bound to it or present in the detection zone and unbound. The intensity plot displaying the protein fluorescence signal as a function of the lipid fluorescent signal presents 3 regions: (i) unbound proteins (single-positive for proteins only in the top left quadrant), (ii) CHMP proteins bound to GUVs (double-positive for proteins and lipids in the top right quadrant), (iii) GUVs free of proteins (single-positive for lipids only in the lower right quadrant).

### QCM-D experiments

Supported lipid bilayers (SLB) were generated with or without PIPs lipids. In the absence of PI(4,5)P2, SLB made of 30% and 40% DOPS-SUV composition were produced with a buffer containing Ca^2+^ (150 mM NaCl, 10 mM Tris (at pH 7.5) + 2 mM Ca^2+^) ^41^. After SLB formation, the bilayer was rinsed with the same buffer but supplemented with EDTA (150 mM NaCl, 10 mM Tris pH 7.5, 10 mM EDTA) to remove Ca^2+^ excess. SLBs were also produced in the presence of PIP lipids (PI(4,5)P2 or PI(3,4,5)P3), with PIP2-SUV or PIP3-SUV lipid compositions, respectively. SLB formation was achieved in a buffer containing 150 mM KCl, 20 mM citrate pH=4.8 ^42^. Following SLB formation, CHMP proteins were injected at concentration of 200 nM in BP buffer. The interaction between the proteins and the lipid bilayer were directly measured from the fifth overtone of the frequency shift (Δϑ_5_).

QCM-D measurements were performed using a Q-Sense E4 system (Q sense; Gothenburg, Sweden). The mass sensor is a silicon dioxide-coated quartz crystal microbalance SiO_2_ (QSX-303 Lot Quantum Design France) with a fundamental frequency of 4.95 MHz. The liquid flow was controlled using a high precision multichannel dispenser (IPC; ISMATEC – Germany). All experiments were performed at room temperature with a flow rate of 50 µL.min^−1^.

### Micropipette experiments

The experimental chamber and the micropipette made of a borosilicate capillary (1 mm outer diameter and 0.58 mm inner diameter (Harvard Apparatus, UK)) introduced into the chamber, are passivated with a β-casein solution at 5 mg.mL^−1^ in Sucrose 25 mM, NaCl 50 mM and Tris 25 mM (pH 7.5) for 15 min. The chamber is rinsed twice with BP buffer. Then, PIP2-GUVs pre-incubated with CHMP proteins are added to the chamber. Once the chamber is sealed with mineral oil, the zero pressure is measured and the aspiration assay can begin by decreasing the water height gradually, thus increasing the applied tension on the vesicle.

The explored tensions for the aspiration experiments with the different CHMP proteins range up to 1.6 mN.m^−1^ (corresponding to the membrane enthalpic regime). The software EZ-C1 was used for acquisition of the confocal images.

At high tension, in the enthalpic regime, an apparent elastic stretching modulus of the membrane χ can be deduced from the linear variation of the fractional excess area 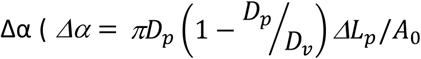 where *ΔL_p_* is the variation of the tongue length and *A*_0_ the initial area of the GUV) as a function of the applied tension σ using 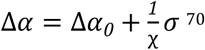, with Δ*α*_0_ being the initial excess area for the reference tension *σ_0_.* According to the Young-Laplace equation, the membrane tension is equal to : 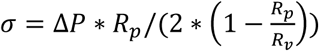 where *ΔP* is the difference of pressure between the interior of the micropipette and the chamber, *R_p_* and *R_v_* are respectively the pipette and vesicle radius ^54^.

### Osmotic shock on GUVs

10% PIP2-GUVs were either co-incubated with 500 nM CHMP2B-ΔC in 50 mM NaCl and 25 mM Tris, at pH 7.4 buffer (CHMP protein binding buffer referred as BP buffer) or transferred to the same buffer free of protein (osmolarity equal to 125 mOsm.L^−1^). CHMP2B-coated GUVs and CHMP2B-free GUVs were then transferred to a hyperosmotic buffer with increasing sodium chloride concentrations up to 250 mM NaCl. The effect of the osmotic shock was visualized using confocal microscopy.

### HS-AFM imaging based deformation experiment

PIP2-SUVs were immobilized on a freshly cleaved mica surface and placed into the AFM chamber with BP buffer. For studying the vesicles with CHMP2B, prior to immobilization to the surface, the PIP2-SUVs were preincubated with 1 µM of CHMP2B for 30 min to obtain full protein coverage on the SUV surface. A high-speed amplitude modulation tapping mode AFM (RIBM, Japan) was used for imaging and deformation experiments, with ultra-short cantilevers (spring constant 0.15 N/m, Nanoworld) ^71^. Initial imaging (at minimum force) was performed at a free cantilever oscillation amplitude of 5.4 nm and a set point amplitude at 4.3 nm. The imaging rate was 1 frame/second. We regulated the set-point amplitude in a stepwise manner, while keeping the free amplitude constant, in order to increase the imaging force. The imaging force can be estimated in first approximation as *F*= *k*Δz, where *k* is the spring constant of the cantilever and *Δz* is the difference between free and set-point amplitude of the cantilever oscillation. It follows that the images were acquired with an estimated minimal force of ∼150 pN. For the measurement of membrane mechanics with and without CHMP2B proteins, image acquisition was first performed at minimal force (∼150 pN). Next step by step the imaging force was increased with 9 % increments, by decreasing the set-point amplitude. After reaching the maximal force, after ∼8 steps and a final imaging force of ∼270 pN, the tapping force was reduced again to its lowest value (150 pN) for each trial. Images were analyzed using IgorPro scripts of the AFM manufacturer (RIBM) and ImageJ scripts. For the determination of the relative spring constant k_rel_ the average slope of all curves in respectively fig. S3E & F is taken and the following transformation was performed k_rel_ = -1/Slope_average_.

## ACKNOWLEDGMENTS.

We acknowledge support from FINOVI (WW), the ANR (ANR-14-CE09-0003-01) (WW, PB), the Institut Universitaire de France (IUF; WW) and from the platforms of the Grenoble Instruct center (ISBG; UMS 3518 CNRS-CEA-UJF-EMBL) supported by the French Infrastructure for Integrated Structural Biology Initiative FRISBI (ANR-10-INSB-05-02) and GRAL (ANR-10-LABX-49-01) within the Grenoble Partnership for Structural Biology (PSB). We furthermore acknowledge funding though a NWO VIDI grant (WHR) and an EU Marie Curie fellowship (INTERACT 751404), (S.Mai.). P.B. group greatly acknowledges the Nikon Imaging Center (Institut Curie, Paris), member of the France BioImaging national research infrastructure (ANR-10-INSB-04) as well as the Flow Cytometry Platform of the Institut Curie for technical support in microscopy and flow cytometry respectively. M.A. was funded by the Université Pierre et Marie Curie/sorbonne Université, Doctoral school « Physique en Ile de France » (ED-564) and the Fondation pour la Recherche Médicale. N.D.F was funded by post-doc fellowships from the Institut Curie, the Fondation pour la Recherche Médicale a EMBO non-stipendiary long term fellowship (ALTF 818-2016) and the European Union’s Horizon 2020 research and innovation program (MSCA No. 751715). P.B. group belongs to the CNRS consortium CellTiss, to the Labex CelTisPhyBio (ANR-11- 798 LABX0038) and to Paris Sciences et Lettres (ANR-10-IDEX-0001-02).

## AUTHOR CONTRIBUTIONS

N.M. performed the expression, purification and labelling of the proteins. M.A. and N.D.F. performed the GUV preparation, the confocal imaging, the FACS and the micropipette experiments. M.A. prepared the SUV experiments for QCM and AFM. M.A. and M.B. performed the QCM-D experiments. S.Mai. performed the AFM experiments. Experiments have been designed and discussed by W.W., W.R., P.B. and S.Man.. M.A., W.W., P.B. and S.Man. wrote the manuscript. The results and their interpretation were discussed by all of the authors.

---

**Supplementary Figure 1: Evolution of the ESCRT-III complex**

**(A)** Table illustrating the ESCRT-III complex function, origin and homologs in *S. Cerevisiae* and *H. Sapiens*.

**(B)** Distribution of Vps2 and Vps24 genes across Eukaryotes showing the presence of two Vps2 genes in high organisms.

**(C)** Table illustrating the implication of ESCRT-III subunits in different subcellular locations in *S. Cerevisiae* and *H. Sapiens*. The names are the human homologues in case of *S. cerevisiae*.

**Supplementary Figure 2: Study of CHMP protein-membrane interaction**

**(A)** Optimization of the buffer conditions to optimize the binding of CHMP2B-ΔC (noted here CHMP2B) at 500 nM. Pre-formed vesicles were incubated with CHMP2B-ΔC in buffers with different salt concentrations ranging from 0 mM to 100 mM NaCl (+Tris 25mM at pH 7.5) and imaged with confocal microscope after 30 min incubation. Lipid signal is shown in magenta and protein signal in green. Scale bar, 5 µm.

**(B)** Confocal image of MBP-CHMP2A-ΔC (noted here CHMP2A) without TEV (Top line) and in the presence of TEV to cleave the MBP tag (Bottom line). Saturated protein fluorescent signal is represented in yellow. Cleavage of MBP tag slightly increases the interaction but induces aggregation. Scale bar: 30 µm

**(C)** Histograms of CHMP2B-ΔC protein fluorescence intensity for PI(3)P, PI(4)P, PI(3,5)P2 and PI(4,5)P2 GUVs (lipid composition 2).

**(D)** Comparison of the binding density of MBP-CHMP2A-ΔC + CHMP3 and of CHMP2B-ΔC to GUVs with different charged lipids, measured by FACS, corresponding to Fig. 1D. The values are normalized to their respective binding density to DOPS. **=p-value<0.01 (Student’s t-test). N = 4 (number of FACS experiment with about 10^4^ counted events per experiment, per condition).

**(E)** QCM-D experiment displaying the typical frequency shift of -25 Hz after supported bilayer formation and a frequency shift Δυ_5_ representative of the amount of protein bound to the bilayer.

**(F)** Spinning disk images of interaction of MBP-CHMP2A-ΔC + CHMP3 in BP buffer on 10% PI(4,5)P2-containing GUVs. CHMP2A-ΔC fluorescent signal is displayed. A z-projection is represented. The different panels corresponding to 3 representative GUVs show the homogeneous coverage of the co-polymer as a function of protein concentration and incubation time. First panel: CHMP2A and CHMP3 are incubated at 500 nM and 2µM, respectively, for 15 min. Second panel: CHMP2A and CHMP3 are incubated at 1 µM and 4µM, respectively, for 15 min. Third panel: CHMP2A and CHMP3 are incubated at 500 nM and 2µM, respectively, for 60 min. Scale bar, 10 µm.

**Supplementary Figure 3: Deformation of vesicles with and without CHMP 2B.**

**(A, B)** Reduction of vesicle height under increasing force for vesicles without and with CHMP2B- ΔC, respectively. ‘Zero’ force increment represents the minimum imaging force (∼150 pN).

**(C)** Example of deformation for a ∼60 nm vesicle over increasing force up to 80% of the initial imaging force.

**(D)** Represents the transformation of vesicle height to relative height for each point for the curve in **C**.

**(E, F)** represent the relative height vs force increment for all the curves from panel **A** and **B**, respectively.

**Supplementary movie S1:** Typical example of vesicle response upon increasing and decreasing imaging force. It can be observed that the vesicle restores its height after the complete decrease in imaging force.

