## Supplementary Figure 1 for "The ESCRT-III Isoforms CHMP2A And CHMP2B Display Different Effects On Membranes Upon Polymerization"

**SUPPLEMENTARY 1**

**A**

| ESCRT complex | Proposed Function | Evolutionary origin | Yeast | Human |
|---|---|---|---|---|
| **ESCRT -III** | Membrane remodelling | Archaea | Vps20<br>Snf7<br>Vps24<br>Vps2 | CHMP6<br>CHMP4A, B, C<br>CHMP3<br>CHMP2A, B |

**B**

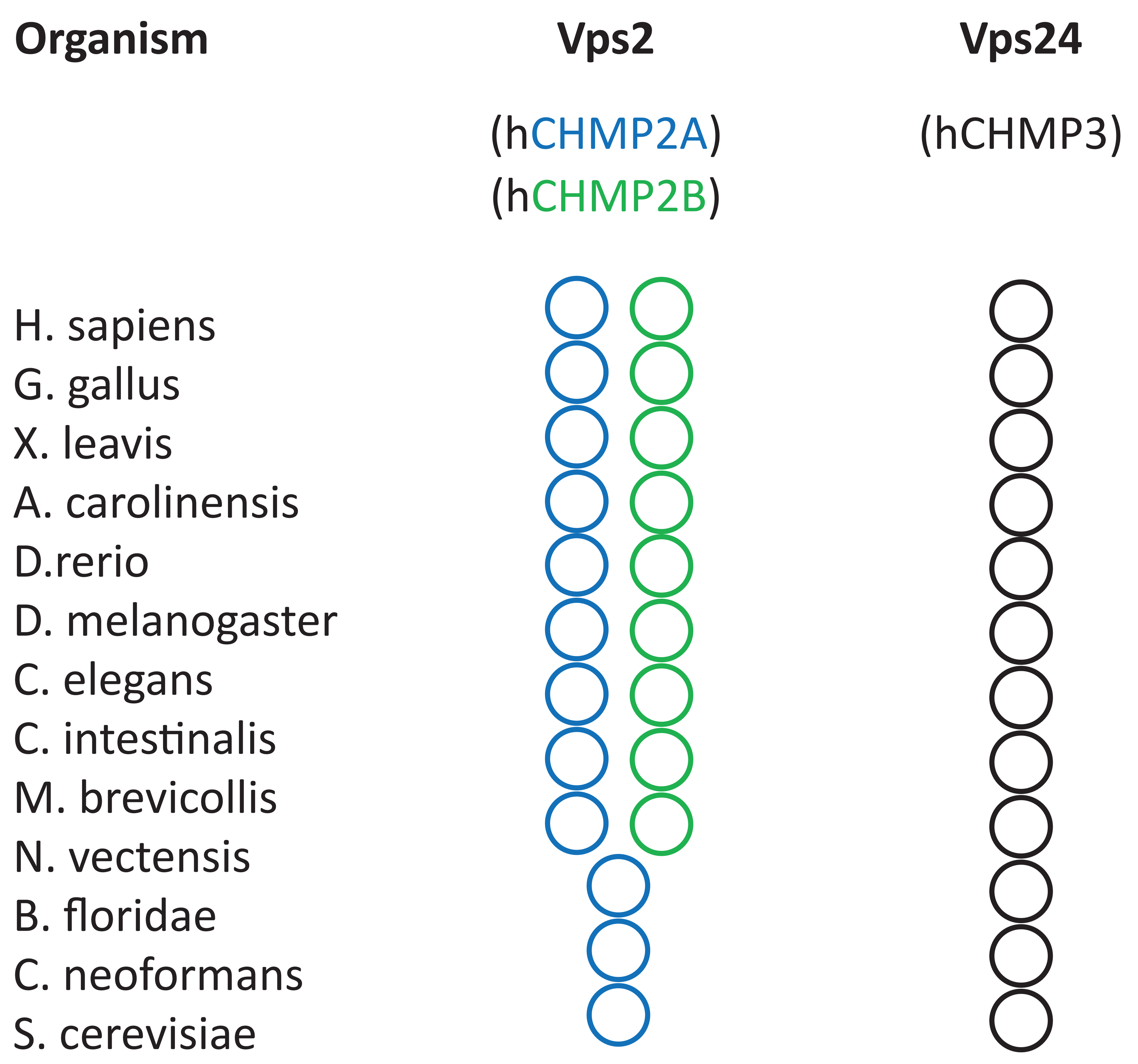


**C**

| Cellular process | Organism | ESCRT -III core components |
|---|---|---|
| **MVB formation** | S. cerevisiae<br>H. sapiens | CHMP4B - CHMP3<br>CHMP2A |
| **Cytokinesis** | H. sapiens | CHMP4B - CHMP3<br>CHMP2A - CHMP2B |
| **HIV-1 budding** | H. sapiens | CHMP4B - CHMP3<br>CHMP2A - CHMP2B |
| **Neurone severing** | H. sapiens | CHMP4B - CHMP3<br>CHMP2A - CHMP2B |
| **Pasma membrane repair** | H. sapiens | CHMP4B - CHMP3<br>CHMP2A - CHMP2B |
