## Supplementary figures and images for "The ESCRT-III Isoforms CHMP2A And CHMP2B Display Different Effects On Membranes Upon Polymerization"

### Supplementary Figure 2

**SUPPLEMENTARY 2**

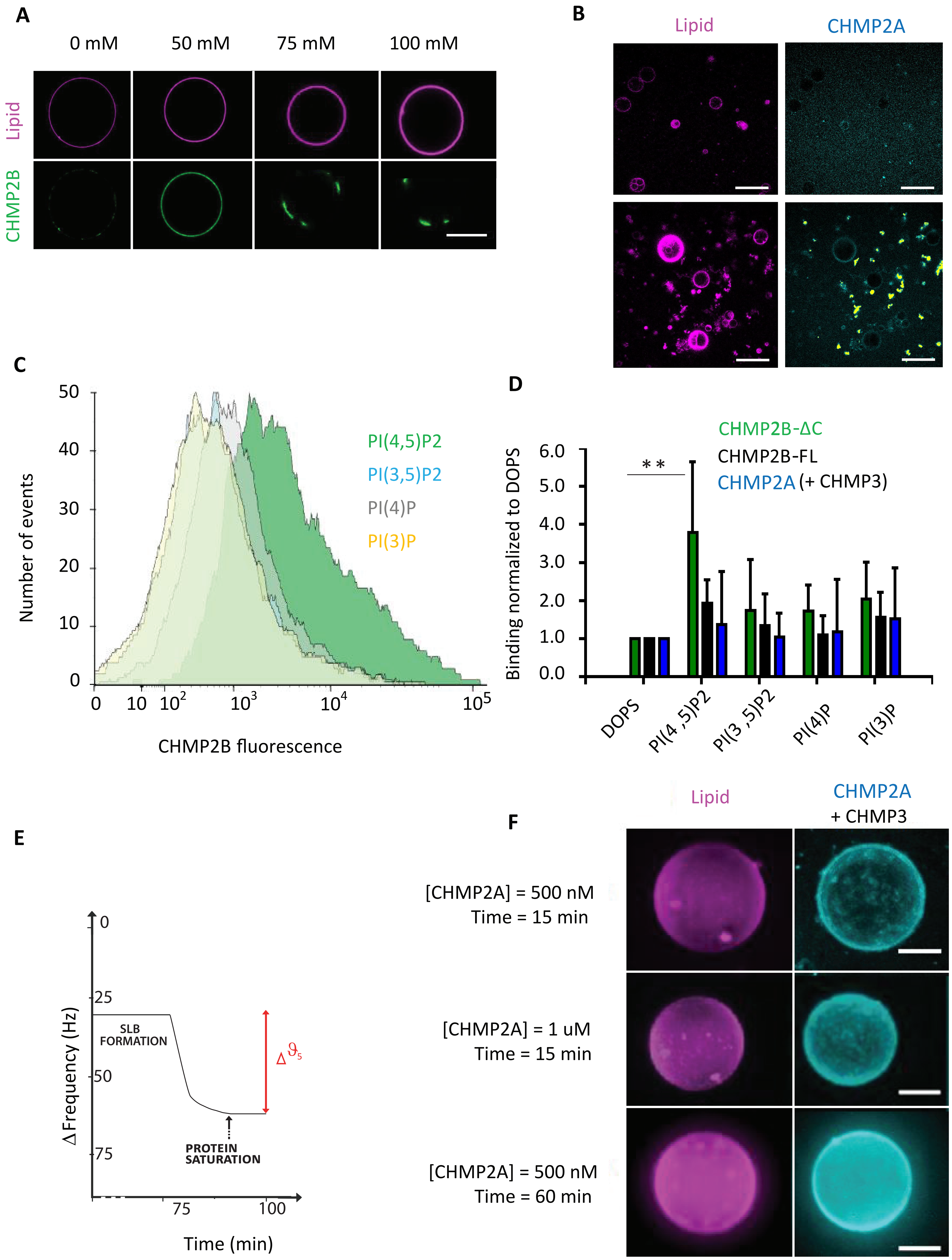

### Supplementary Figure 3

**SUPPLEMENTARY 3**

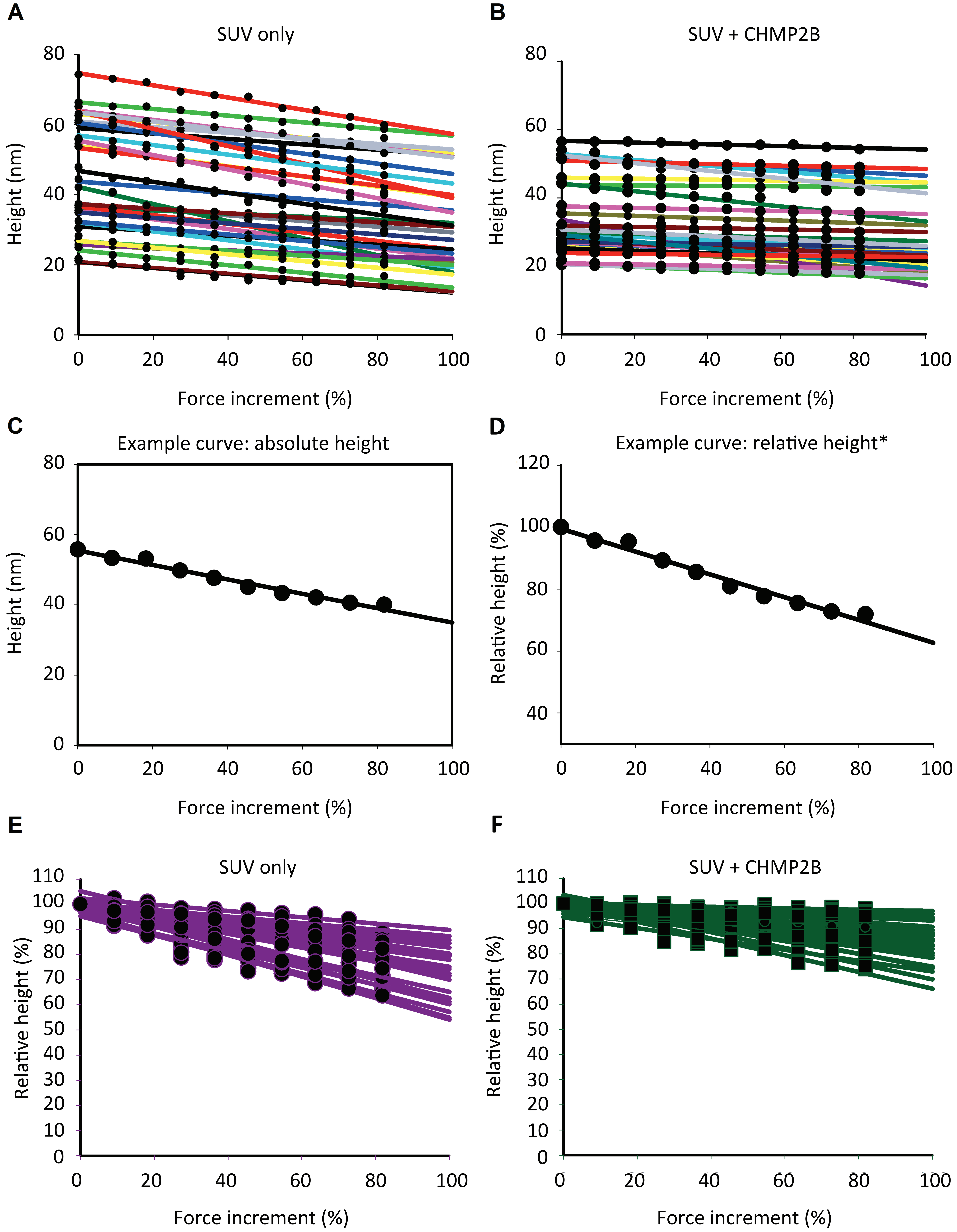
